# A horizontally acquired cyclic di-GMP phosphodiesterase regulated by zinc and quorum sensing

**DOI:** 10.1101/2025.04.25.650742

**Authors:** Aathmaja Anandhi Rangarajan, Kiwon Ok, Marissa K. Malleck, Micah J. Ferrell, Thomas V. OˈHalloran, Christopher M. Waters

**Affiliations:** Department of Microbiology Genetics and Immunology, Michigan State University, East Lansing, MI, USA; Elemental Health Institute, Michigan State University, East Lansing, MI, USA; Department of Chemistry, Michigan State University, East Lansing, MI, USA

## Abstract

The second messenger cyclic di-GMP (cdG) in *V. cholerae* is indispensable for the regulation of biofilm formation, motility, and a variety of other important bacterial behaviors in the majority of bacteria. The human pathogen *Vibrio cholerae* has a diverse repertoire of diguanylate cyclase (DGC) and phosphodiesterase (PDE) enzymes that control the intracellular cdG concentration depending on local environmental cues and its physiological state. Determining the transcriptional regulation of these enzymes and the respective environmental signals that control their activity is important to understand how and when *V. cholerae* switches between motile and sessile lifestyles in different environments. In some strains of the current *V. cholerae* 7^th^ pandemic El Tor biotype, the horizontally acquired *Vibrio* Seventh Pandemic 2 (VSP-2) island encodes an uncharacterized PDE at the gene locus *vc0515* which we named *zpdA* (**Z**inc-inhibited **P**hospho**d**iesterase-**A**). We show here that *zpdA* transcription is repressed by Zur when Zn^2+^ is abundant, as well as by the quorum sensing regulator HapR when cells grow to high density. Furthermore, we find that the PDE activity of the purified ZpdA protein is inhibited by Zn^2+^ but is dependent on alternative divalent cations such as Mn^2+^, which we find is elevated in *V. cholerae* cells grown under zinc limiting conditions. We conclude that ZpdA is an active metal-dependent PDE that is regulated by Zn^2+^ availability at both the level of transcription and post-translation leading to elevated cdG levels when Zn^2+^ is abundant. Our results demonstrate the important role of metal availability in modulating cdG signaling in bacteria.

**Importance:** *V. cholerae* colonizes estuarine environments and human host where it transitions between motile to sessile states which are controlled by cdG levels. cdG levels change in response to a variety of signals and are controlled by the activity of DGCs and PDEs, enzymes that make and degrade cdG. In this work, we show that Zn^2+^ and the cell density regulator HapR repress ZpdA, the PDE present in the VSP-2 island, at the level of transcription, and that Zn^2+^unexpectedly alters the PDE activity of ZpdA protein itself. Our study highlights the role of metal availability as an important signaling cue that controls *V. cholerae* biology.

## Introduction

*V. cholerae* is a bacterial pathogen capable of causing cholera. One distinguishing feature of the El Tor biotypes, which are responsible for the 7^th^ and current pandemic of *V. cholerae,* from the previous classical biotype is the acquisition of the Vibrio Seventh Pandemic Islands 1 (VSP-1) and -2 (VSP-2). These islands are hypothesized to have aided El Tor in displacing the Classical biotype in both environmental and clinical reservoirs^1,2^. Although many of the gene functions in these genomic islands are unknown, some of the genes of VSP-1 and VSP-2 islands are involved in defense against phages and foreign DNA^3–6^. In addition to infecting humans, *V. cholerae* associates with various species in aquatic environments where it is challenged with diverse stressors. Thus, strategies to successfully switch between motile and sessile forms in these environments are also likely key to the emergence of El Tor^7^.

The second messenger cdG, controls several important physiological processes, including but not limited to biofilm formation, motility, type two secretion, DNA repair, cell shape, and colonization^8–12^. The intracellular cdG concentration is a major driver of the motile-sessile lifestyle switch whereby high levels of cdG promote biofilm formation and low levels induce motility^10^. Intracellular cdG concentrations are dictated by the regulation of GGDEF containing diguanylate cyclases (DGCs) which synthesize cdG and EAL/HD-GYP containing phosphodiesterases (PDEs) which degrade it ^8,9^. The first sequenced strain of *V. cholerae* possess 31 GGDEF containing proteins and 10 that contains both an EAL and GGDEF, 12 that contains EAL and 8 that contain HD-GYP domains, although such numbers can exhibit minor variation from strain to strain^8,10^. For example, one of the PDE containing proteins, *vc0515*, which we rename in this work to *zpdA* for **<u>Z</u>**inc-inhibited **<u>P</u>**hospho**<u>d</u>**iesterase-**<u>A</u>**, is encoded in the VSP-2 island in only some strains of El Tor, suggesting a horizontally acquired PDE can potentially impact cdG levels and thereby regulate biofilm formation and motility^13^. However, the contribution of ZpdA to cdG, biofilm formation, and motility nor the environmental signals that regulate this protein are not well understood.

Most cdG DGCs or PDEs possess one or more N-terminal sensory domains that are thought to assimilate and respond to specific signals^8,14^. However, only a few signals, such as nitric oxide, bile, spermidine, light, and oxygen, have been described to regulate specific DGC and PDE activity in *V. cholerae,* thereby modulating intracellular cdG^14,15^. Metals such as Zn^2+^, Fe^3+^, Mn^2+^, and Mg^2+^, are essential nutrients, which unlike other organic enzymatic co-factors, can neither be synthesized nor destroyed. Thus the intracellular concentrations of these nutrients are maintained in narrow ranges through tightly regulated homeostasis mechanisms under high and low metal conditions^16–18^. Mn^2+^, Ca^2+^, and Fe^3+^ were shown to regulate *in vitro* activity of VieA, *Vc*EAL, and VCA0681, respectively in *V. cholerae*^19–21^. Besides these *in vitro* studies, whether and how these metals impact cdG regulation in *V. cholerae* or more broadly, in other bacterial species is not understood.

Here, we show that the *zpdA* PDE allows *V. cholerae* to regulate cdG levels in response to Zn^2+^ availability. This occurs both transcriptionally and by direct regulation of ZpdA activity. At high intracellular Zn^2+^ concentrations, the *vc0513-15* operon is repressed by the Zn^2+^-responsive repressor Zur. Low Zn^2+^ levels lead to *zpdA* expression via Zur de-repression. We also found that Zn^2+^ directly inhibits the PDE activity of ZpdA, demonstrating that increases in Zn^2+^ availability in the cell leads to elevation of cdG concentrations through both transcriptional and posttranscriptional processes. We also demonstrate that the transcription of *zpdA* itself, but not the *vc0513-15* operon, is additionally repressed by the quorum sensing master regulator HapR. The *vc0513-15* operon is highly conserved in several El Tor strains within the highly variable VSP-2 region, underlining the potential role of Zn^2+^ modulation of cdG in the environmental persistence of the 7^th^ *V. cholerae* pandemic.

## Results

### ZpdA is an active PDE encoded in the VSP-2 island

The *zpdA* (*vc0515*) gene encodes for a predicted EAL PDE. This gene is located in the *vc0513-15* operon of the VSP-2 island of some El Tor strains, including N16961, and this operon was previously demonstrated to be repressed by the Zn^2+^-responsive repressor Zur^13^ (Fig. 1A). Unlike most PDEs, the EAL domain of ZpdA is encoded on the N-terminus of the protein, while the C-terminus encodes an HDOD domain, which has been shown to be regulated by metals^22,23^. We therefore sought to determine the impact of ZpdA, and its modulation by Zur and Zn^2+^, on intracellular cdG and associated phenotypes in *V. cholerae* strain N16961.

**Fig. 1:**
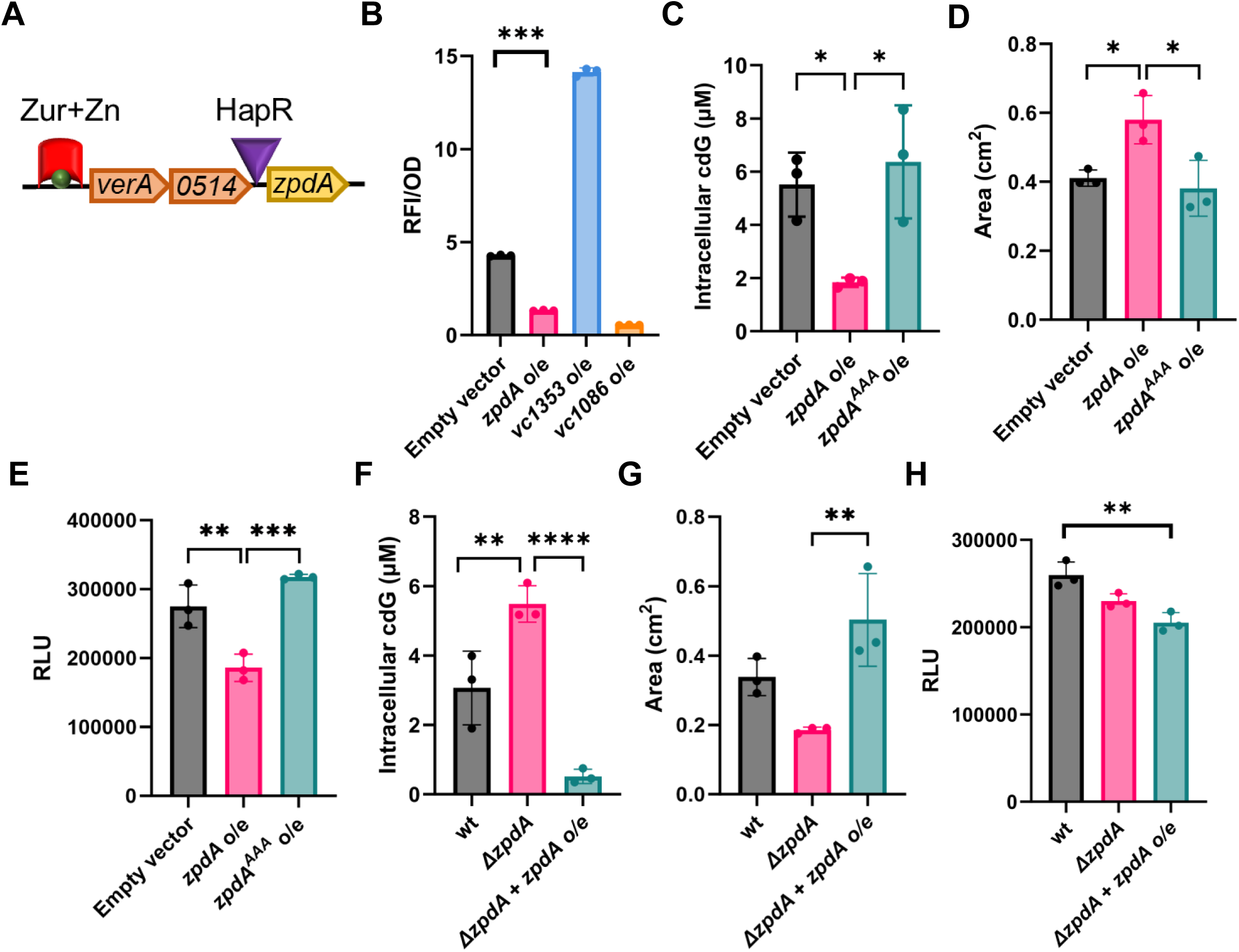
ZpdA is an active phosphodiesterase(A) Genomic loci map of *verA, vc0514* and *zpdA* genes. Zur repressor (red) forms complex with divalent zinc ions (green) and binds to *verA* promoter. Potential HapR binding site (purple) upstream of *zpdA* coding region is shown. (B) CdG binding RFP biosensor readout normalized to OD from overexpression of DGC (*vc1353*), PDE (*vc1086*), PDE (*zpdA*) and empty vector (pEVS141) with 1 mM IPTG. (C) Intracellular cdG levels in strains overexpressing *zpdA*, *zpdA^AAA^* or empty vector (pEVS141) with 1 mM IPTG. (D) Quantification of motility by measuring the colony area in strains overexpressing *zpdA*, *zpdA^AAA^* or empty vector (pEVS141) with 1 mM IPTG. (E) Quantification of biofilm expressed as relative luminescence units in strains overexpressing *zpdA*, *zpdA^AAA^* or empty vector (pEVS141) with 1 mM IPTG. (F) Intracellular cdG levels in wild-type, *ΔzpdA* and *ΔzpdA* with overexpression of *zpdA.* (G) Quantification of motility by measuring the colony area in strains in wild-type, *ΔzpdA* and *ΔzpdA* with overexpression of *zpdA.* (H) Quantification of biofilm expressed as relative luminescence units in wild-type, *ΔzpdA* and *ΔzpdA* with overexpression of *zpdA.* The mean and standard deviation of three biological replicates is indicated. Statistical significance was determined by one-way ANOVA and post-hoc Tukey test (* = p < 0.05, *** = p < 0.005, **** = p < 0.0005)

To determine if ZpdA is an active PDE, we overexpressed *zpdA* along with a cdG biosensor (pRP0122-*Pbe_amcyan_Bc3-4_turborfp*) in minimal media. This biosensor contains three cdG-responsive riboswitches that control the expression of *turboRFP* such that increased intracellular cdG leads to increased RFP^24^. *zpdA* overexpression decreased RFP expression similar to the PDE *vc1086,* which was previously shown to be active in these conditions, indicating a decrease in cdG^25^. In contrast, expression of the DGC *vc1353* increased RFP levels, reflecting increased cdG (Fig. 1B). ZpdA possesses the amino acids “ELL” in its active site, and indeed overexpression of the active site mutant (ELL-AAA) of *zpdA*, which we name *zpdA^AAA^*, followed by direct quantification of intracellular cdG using LC-MS/MS, indicated that WT *zpdA* reduced cdG while this mutation rendered it inactive (Fig. 1C). Consistent with its impact on intracellular cdG, overexpression of *zpdA* increased motility and decreased biofilm formation when compared to the empty vector or the *zpdA^AAA^*(Figs. 1D, 1E).

To determine whether *zpdA* impacted cdG at native expression levels, we measured the intracellular cdG levels in the WT strain and an isogenic *ΔzpdA* mutant in M9 minimal media. The *ΔzpdA* mutation exhibited higher intracellular cdG compared to the WT strain, which could be offset by expression of *zpdA* from a plasmid (Fig. 1F). The elevated intracellular levels of cdG in the *ΔzpdA* mutant led to decreased motility (Fig. 1G). However, the *ΔzpdA* mutant did not significantly impact biofilm formation relative to the wild-type strain (Fig. 1H). Importantly, measuring cdG, motility, and biofilms requires different growth conditions that might impact the activity or expression of ZpdA. Moreover, given that *V. cholerae* encodes dozens of PDEs and cdG concentrations can be modulated via negative feedback loops, it is common not to observe a significant phenotypic difference when only one is deleted^25^. The summation of these results indicates that *zpdA* is an active PDE in the conditions examined here that decreases intracellular cdG, impacting downstream cdG-regulated phenotypes.

### Zn^2+^ inhibits the expression of *zpdA*

Transcription of the *zpdA* gene is repressed by the Zur repressor in the presence of elevated Zn^2+^ availability through the upstream *verA* (*vc0513*) promoter^13^ (Fig. 1A). Deletion of the *zur* gene did not significantly impact intracellular cdG and motility while leading to decreased biofilm formation (Fig. S1A-C). Decreased biofilm formation in the *zur* mutant could be due to the downregulation of several *rbm* and *vps* operon genes involved in biofilm formation, as previously demonstrated in a transcriptomic analysis^13^.

Given that interpretation of the Δ*zur* mutant is challenging, likely because it regulates multiple genes^13^, we analyzed the impact of Zn^2+^ on *zpdA* expression. First, we sought to understand how zinc levels changed in different growth conditions and after adding extracellular Zn^2+^. We measured the level of zinc in different media powders and final growth media using Inductively Coupled Plasma Mass Spectrometry (ICP-MS). As a control, we added 10 µM of Zn^2+^ to minimal media and measured it alongside other media. Minimal media, minimal media with zinc, and LB media contained 0.03 µM, 11 µM, and 15 µM of zinc, respectively (Fig. 2A). *Vc* colonizes crustaceans as biofilms and utilizes chitin as a carbon source, hence we also measured the level of zinc in chitin^26,27^. In the laboratory, *Vc* is grown with chitin flakes in Instant Ocean Water medium resembling the ocean water as described in the methods. Therefore, we analyzed the zinc content of chitin flakes from shrimp and crab alongside instant ocean salts. Shrimp chitin (0.0014 ng/µg) and crab chitin (0.06 ng/µg) had significantly higher zinc content than instant ocean salts alone (0.00013 ng/µg) (Fig. S2).

**Fig. 2:**
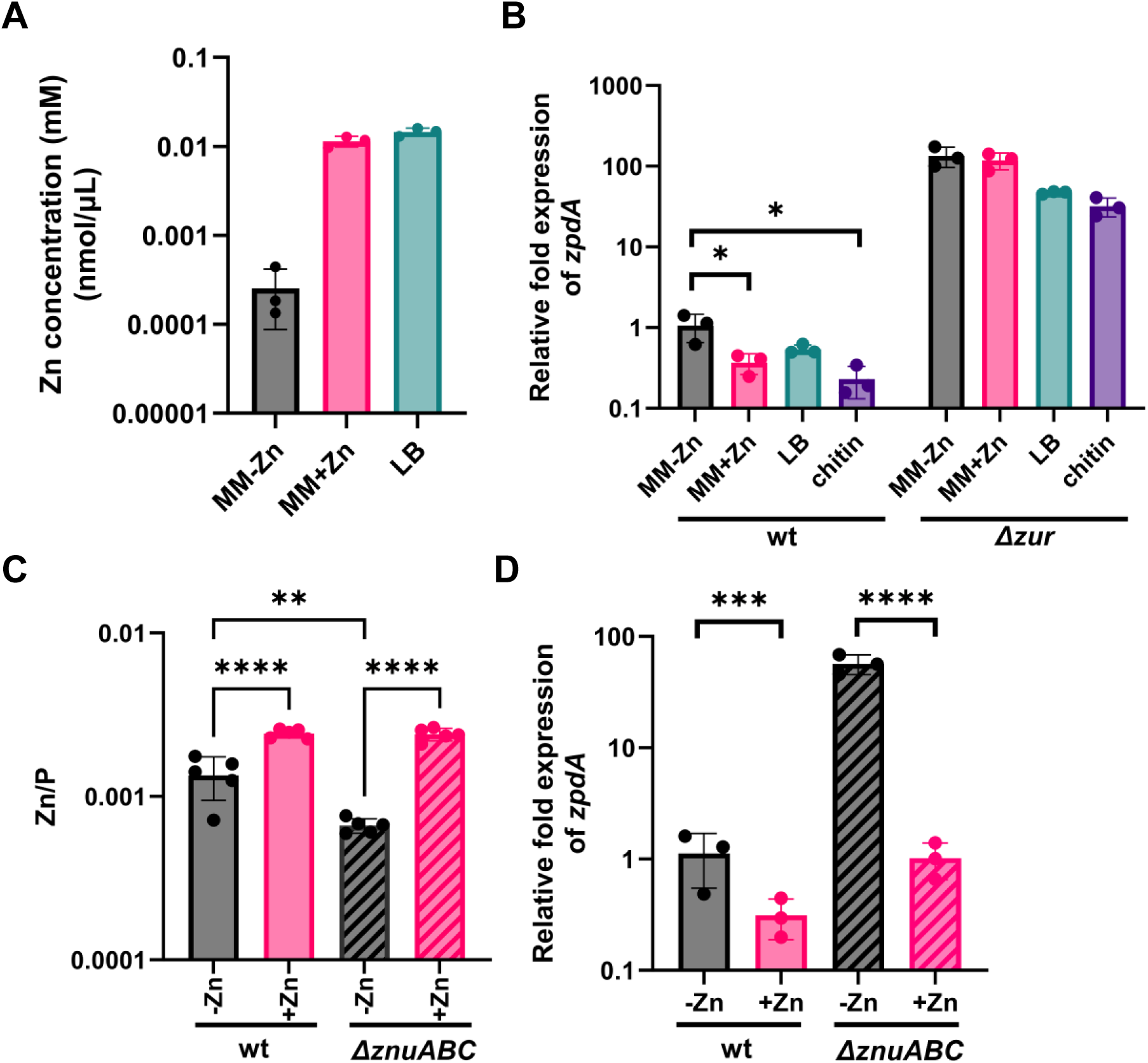
Zur in the presence of Zn^2+^ represses expression of the *zpdA* phosphodiesterase (A) Inductively Coupled Plasma Mass Spectrometry (ICP-MS) quantification of zinc content in minimal media, minimal media with zinc sulfate (10 µM), and LB media. Average of three independent replicates is indicated. (B) Relative expression of *zpdA* determined by qPCR in wild-type and *Δzur* in minimal media, minimal media with zinc, chitin and LB (C) ICP-MS quantification of intracellular zinc normalized to phosphorus in wild-type and *ΔznuABC* mutant in minimal media with and without zinc supplementation (10 μM ZnSO_4_). (D) Relative expression of *zpdA* determined by qPCR in wild-type and *ΔznuABC* in minimal media and minimal media plus zinc. Mean and standard deviation of three biological replicates is indicated. Statistical significance was determined by one-way ANOVA and post-hoc Tukey test (* = p < 0.05, *** = p < 0.005, **** = p < 0.0005)

With knowledge of zinc concentrations in these media, we next determined the relative expression of *zpdA* in different media in the WT strain and *Δzur* mutant using qRT-PCR. In the WT strain, the relative expression of *zpdA* was decreased 2-5-fold in LB, crab chitin, and minimal media with zinc supplementation (10 μM) when compared to minimal media alone (Fig. 2B). We observed a more dramatic effect in the *Δzur* strain where expression of *zpdA* was 88-325 fold higher and was non-responsive to zinc addition in minimal media (Fig. 2B). This result clearly shows that Zur is an active repressor in all media examined, which is likely due to the sub-femtomolar affinity of Zur for zinc leading to repression even at low zinc concentrations^17^.

A previous study showed de-repression of upstream *verA* promoter using the zinc importer *ΔznuABC* mutant^28^. We therefore reasoned that this mutant could be used to assess the effects of zinc on *zpdA* repression, given it may have lower intracellular concentrations of zinc. To confirm this, we quantified the intracellular concentrations of zinc and normalized it to phosphorus levels in each sample, which is an established internal standard for cell biomass using ICP-MS^29^. Normalized zinc levels were 2-fold lower in the *ΔznuABC* mutant relative to wild-type. Adding 10 μM ZnSO_4_ to minimal media increased the intracellular zinc levels in both wild-type and *ΔznuABC* strains (Fig. 2C). High zinc concentrations in the *ΔznuABC* mutant upon zinc addition is likely due to secondary zinc transporters^30^. In line with the intracellular zinc levels, the relative expression of the *zpdA* gene was 57-fold higher in *the ΔznuABC* mutant when compared to the WT, and the addition of zinc strongly repressed *zpdA* expression (Fig. 2D). To assess the impact of ZnuABC on the import of other metals, we also measured other metal ions in WT and *ΔznuABC* with and without additional zinc using ICP-MS. There was no significant difference between the WT and *ΔznuABC* mutant for Mn, Mg, Na, K, Fe, Ni, Ca, Cu and Cr contents (Fig. 3). However, we did observe a statistically significant reduction of Cu levels in the *ΔznuABC* mutant of 1.7-fold (Fig. 3). Zinc addition did not have any significant impact on metal concentrations except for Mn levels, which decreased 3.5-fold upon the addition of ZnSO_4_ to WT cells. Zinc impacts on Mn levels also occurred in the Δ*znuABC* mutant, demonstrating these differences are not due to this zinc transporter (Fig. 3). These results demonstrate that changes in the intracellular concentration of zinc modulate Zur repression of *zpdA*, but normally, laboratory growth conditions have sufficient intracellular Zn^2+^ for it to function as a Zur corepressor.

**Fig. 3:**
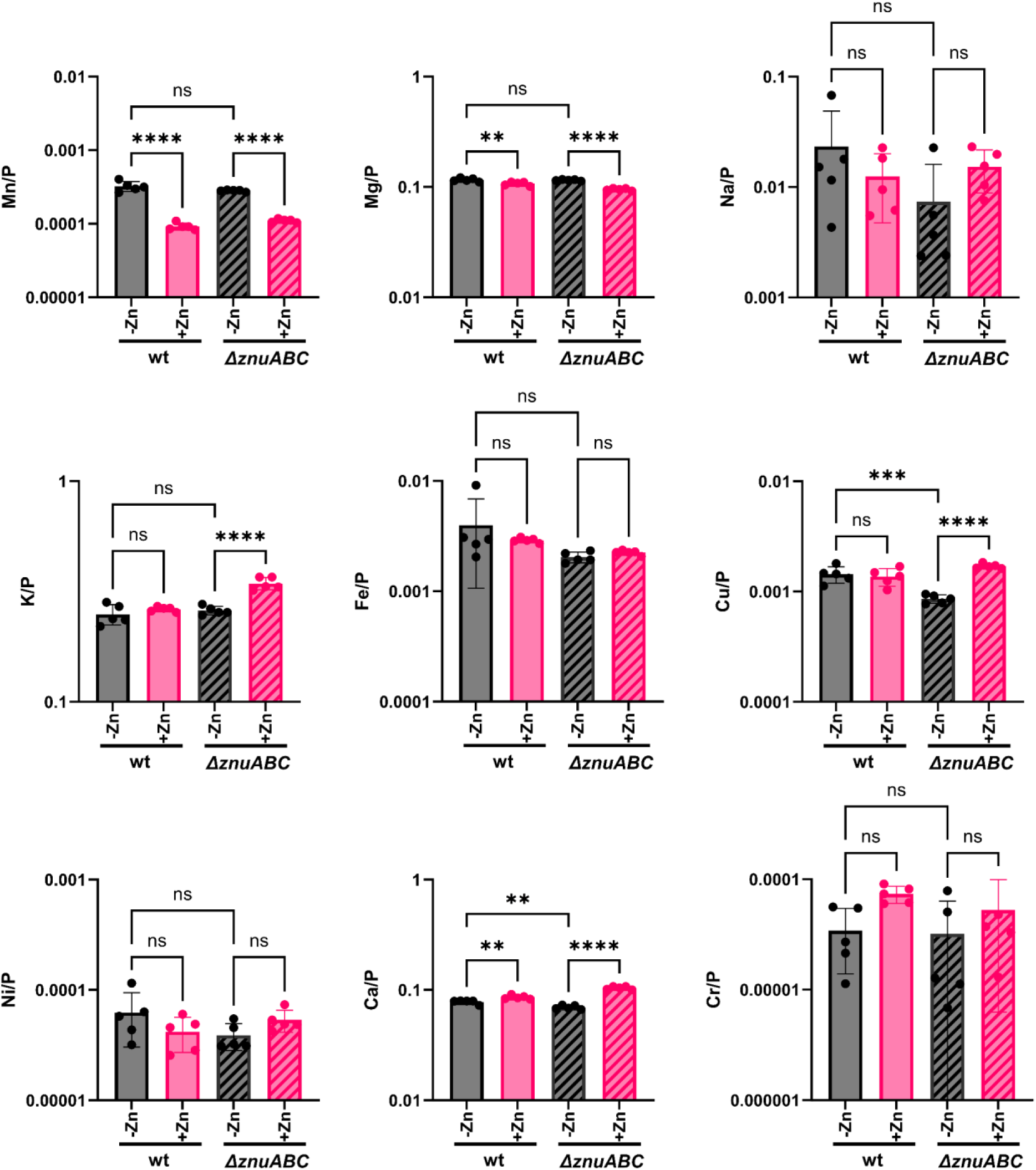
Intracellular metal concentrations of Mn, Mg, Na, K, Fe, Ni, Ca, Cu and Cr (note, different charge forms cannot be discriminated by ICP-MS) content normalized to phosphorus atoms in wild-type and the *ΔznuABC* mutant grown in minimal media and minimal media supplemented with 10 µM ZnSO_4_ determined by ICP-MS. Mean and standard deviation of three biological replicates is indicated. Statistical significance was determined by one-way ANOVA and post-hoc Tukey test (* = p < 0.05, *** = p < 0.005, **** = p < 0.0005).

### Repression of *zpdA* by the quorum sensing (QS) regulator HapR

*zpdA* is part of an operon of genes containing *verA, vc0514,* and *zpdA* encoding a promoter upstream of *verA* (Fig. 1A), and it is thought to be expressed as a single gene transcript via its own promoter^28^. However, bioinformatic analysis predicted a HapR binding site directly upstream of the *zpdA* coding region^30^. HapR is the master quorum sensing (QS) regulator in *V. cholerae* that is expressed when cells transition to high cell density, regulating hundreds of genes including those coding for DGCs and PDEs^30,31^. To determine if *zpdA* is directly regulated by HapR, we generated a plasmid encoded transcriptional fusion of the 100 bp upstream of *zpdA* containing the putative HapR binding site to *luxCDABE*. Since *V. cholerae* strain N16961 encodes a non-functional *hapR*, which has been observed for many El Tor strains, we analyzed expression of *P_zpdA_-lux* expression at the low and high cell density states using the QS-competent *V. cholerae* strain C6706. We also examined a *ΔluxO* mutant (locking the cells at high cell density) and a *ΔhapR* mutant (locking the cells at low cell density). Each of these strains also encoded a deletion of *vpsL* to abolish biofilm formation, enabling accurate readout of the transcriptional reporter. P_zpdA_*-lux* expression was higher in the QS WT strain (Δ*vpsL*) at low cell density when compared to the high cell density condition where HapR is expressed (Fig. 4A). Similarly, the expression was constitutively high in the Δ*hapR* Δ*vpsL* mutant and repressed in Δ*luxO* Δ*vpsL* where HapR is constitutively expressed (Fig. 4A). Overexpression of HapR in strain N16961 had a similar effect driving P_zpdA_*-*lux repression (Fig. 4B). These results combined with the predicted HapR binding site suggest that HapR can bind to *zpdA* upstream region to repress its expression.

**Fig. 4:**
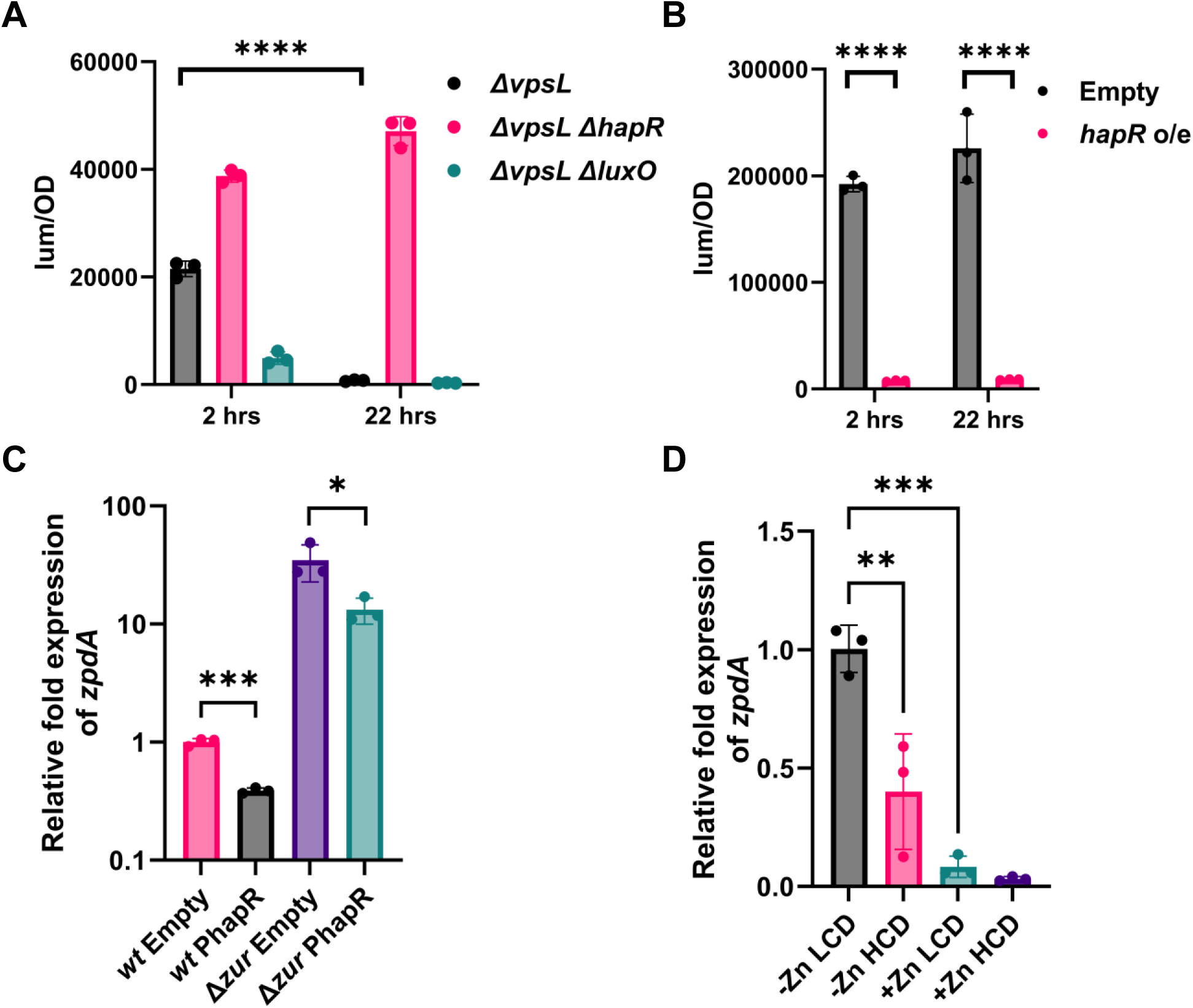
HapR represses the *zpdA* promoter region (A) Luminescence measured from the expression of *P_zpdA_-lux* normalized to OD_600_ in *ΔvpsL*, *ΔvpsL ΔluxO*, *ΔvpsL ΔhapR* strains of C6706 at 2 hours and 22 hours. (B) Luminescence measured from the overexpression of HapR and empty vector normalized to OD_600_ in *ΔvpsL* strain of N16961. (C) Relative expression of *zpdA* determined by qPCR in wild-type and *Δzur* strains of N16961 with overexpression of HapR or empty vector. (D) Relative expression of *zpdA* in E7646 strain at low cell density (LCD) (OD – 0.1) and high cell density (HCD) (OD_600_ 1.5-2) in minimal media with and without Zn^2+^. Mean and standard deviation of three biological replicates is indicated. Statistical significance was determined by one-way ANOVA and post-hoc Tukey test (* = p < 0.05, *** = p < 0.005, **** = p < 0.0005)

Our results demonstrate that both Zur and HapR repress expression of *zpdA.* To disentangle the impact of each of these regulators, we tested the relative expression of *zpdA* using qRT-PCR upon overexpression of HapR in WT N16961 and the *Δzur* mutant. HapR overexpression reduced the expression of *zpdA* 2.6-fold in WT and *Δzur* mutant (Fig. 4C). However, deletion of *zur* alone increased expression of *zpdA* 35-fold, suggesting that in strain N16961 under the conditions tested here, Zur had a larger impact on *zpdA* expression compared with HapR.

We next wanted to test the relative impact of Zn^2+^ and HapR on the expression of *zpdA* under native conditions without HapR overexpression. As N16961 does not encode a functional *hapR* and C6706 does not encode the *zpdA* gene, we performed this experiment in strain E7646, which encodes both a functional *hapR* and the *vc0513-15* operon in the VSP-2 island. We performed this experiment in M9 minimal media with and without Zn^2+^ at low and high cell densities. The addition of Zn^2+^ to this strain significantly decreased expression of *zpdA,* but QS regulation was evident in both conditions as the relative expression of *zpdA* was reduced 2.5- and 2.6-fold between low and high cell density in low and high Zn^2+^ (Fig. 4D). Similar effects of QS in strain E7646 grown on chitin flakes and in LB medium were also observed (Fig. S3A). These results demonstrate that both Zur and HapR regulate the expression of *zpdA,* with Zur playing the dominant role while HapR modulates expression in the conditions tested.

### Zn^2+^ inhibits the PDE activity of ZpdA

cdG signaling allows bacteria to sense environmental changes to regulate biofilm formation and motility. As demonstrated above, Zn^2+^ represses *zpdA* transcription; however, given that the already expressed ZpdA enzyme would remain for some time, such a transcriptional response would exhibit significant phenotypic lag before cdG levels were impacted. We thus considered if Zn^2+^ impacted ZpdA in a post-transcriptional manner, leading to a more rapid response of cdG levels to changing Zn^2+^ concentrations.

In support of this idea, when we overexpressed *zpdA* from a plasmid in strain N16961, we found that ZpdA was active and decreased cdG levels in M9 minimal media but not in LB or minimal media supplemented with Zn^2+^ as measured by direct quantification of cdG (Fig. 5A) or the cdG biosensor (Fig. S3B). Moreover, *ΔzpdA* only exhibited a significant difference in the intracellular levels of cdG compared to the WT strain when grown in minimal media but not in LB or Zn^2+^-supplemented minimal media (Fig. 5B). As LB media is rich in Zn^2+^, these results suggested that Zn^2+^ may directly inhibit the PDE activity of ZpdA.

**Fig. 5:**
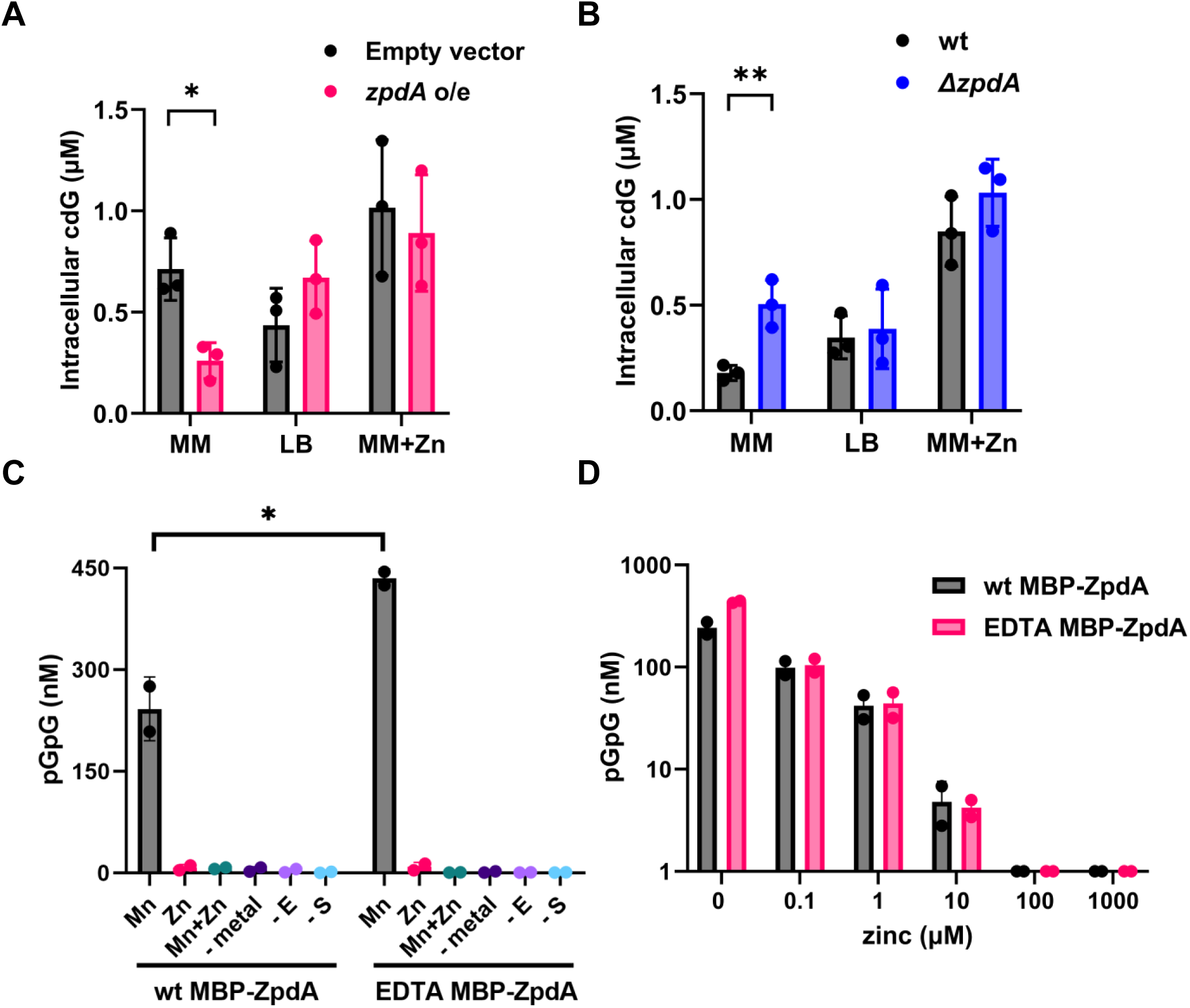
Zn^2+^ inhibits the protein activity of ZpdA (A) Intracellular cdG levels with overexpression of *zpdA* and empty vector in minimal media and LB. (B) Intracellular cdG levels of wild-type and *ΔzpdA* in minimal media and LB. (C) Concentration of pGpG measured by LC-MS/MS in wild-type and EDTA-treated protein with cdG substrate in the presence of 1 mM MnCl_2_, 1mM ZnCl_2_ and 1mM MnCl_2_+ZnCl_2_ along with no metal, no substrate and no enzyme controls. (D) Concentration of pGpG produced from cdG measured by LC-MS/MS in samples of the as-purified protein and EDTA-treated protein with 1 mM Mn and different concentrations of Zn^2+^ (0, 0.1, 1, 10, 100 and 1000 µM). Mean and standard deviation of three replicates is indicated. Statistical significance was determined by t-test (* = p < 0.05, *** = p < 0.005, **** = p < 0.0005).

To test whether Zn^2+^ inhibits the activity of ZpdA, we purified an MBP-ZpdA fusion protein in the presence and absence of EDTA. We used EDTA treatment to remove any inhibitory metals that might bind to ZpdA. We tested the PDE activity of EDTA-treated and untreated MBP-ZpdA using cdG as substrate by measuring the accumulation of 5’-pGpG-3’ using mass LC-MS/MS. Interestingly, without any metal addition, ZpdA was inactive (Fig. 5C).

While the mechanisms of manganese uptake are not yet clear, the increased Mn content under zinc-limiting conditions led us to test for a biochemical role for this divalent metal. We found that MBP-ZpdA exhibited PDE activity only when 1 mM MnCl_2_ or to a lesser extent 1 mM MgCl_2_ was added to the reaction (Fig. 5C and S3C). Addition of 1 mM ZnCl_2_, FeSO_4_ or CaCl_2_ did not stimulate PDE activity, nor did addition of combination of MnCl_2_ and ZnCl_2_ (Fig. 5C and S3C). This result led us to test whether Zn^2+^ was inhibiting PDE activity by conducting a dose response assay. In the presence of 1 mM MnCl_2_, we find that the PDE activity of MBP-ZpdA was decreased in a dose-dependent manner by increasing concentrations of Zn^2+^ (Fig. 5D). Additionally, we find that the Mn^2+^ induced PDE activity was persistently higher for MBP-ZpdA samples that were purified in the presence of EDTA vs the absence of EDTA (Fig. 5C,D). The most likely explanation is that EDTA removed any contaminating Zn^2+^ allowing for higher levels of Mn^2+^ loading and thus leading to higher levels of enzyme activity. Taken together, these experiments demonstrate that Zn^2+^ inhibits the PDE activity of ZpdA protein whereas manganese and, to a lesser extent, magnesium, activates it. This finding aligns with our *in vivo* experiments, which show that ZpdA remains inactive at high Zn^2+^ concentrations.

## Discussion

Although the presence of the VSP-1 and -2 islands are hallmarks of the current *V. cholerae* El Tor biotype, multiple variants of VSP-2 have been identified^2,32^. Our analysis and others have revealed the *vc0513-15* genes in the VSP-2 island genes are highly conserved among several El Tor isolates tested (Fig. S4)^32,33^. For example, a recent isolate responsible for a major outbreak in Bangladesh (BD-1.2) lacks genes *vc0495-vc0512* in the VSP-2 island but encodes *vc0513-15* (Fig. S4)^34,35^. Conservation of these genes in most El Tor isolates underlines their evolutionary significance and potential role in environmental persistence.

The QS regulator HapR has been shown to decrease global cdG levels by transcriptional regulation of DGCs and PDEs^31^. This study was performed in strain C6706, which lacks *vc0513-15* (Fig. S4), and thus the impact of *zpdA* regulation by QS was not assessed. Here, we demonstrate that HapR represses the *zpdA* promoter, although the magnitude of HapR repression on the expression of the *zpdA* gene is minimal when compared to Zur repression from the upstream *vc0513* promoter (Fig. 4C). Although Zur tightly binds to the upstream *verA* promoter and represses *zpdA*, LCD might allow partial de-repression of *zpdA*. N16961 strain with non-functional HapR, grown in minimal media, results in active ZpdA albeit strongly repressed by Zur (Fig. 1F). Thus, QS provides additional regulation of *zpdA*, beyond Zur-mediated repression (Fig. 6). It is also known that several El Tor strains have null mutations in the *hapR* gene, resulting in elevated intracellular cdG^36,37^. Hence, the acquisition of *zpdA* in the VSP-2 island might reduce cdG levels in these strains, thereby enabling cdG homeostasis to effectively switch between motile and sessile phenotypes.

**Fig. 6:**
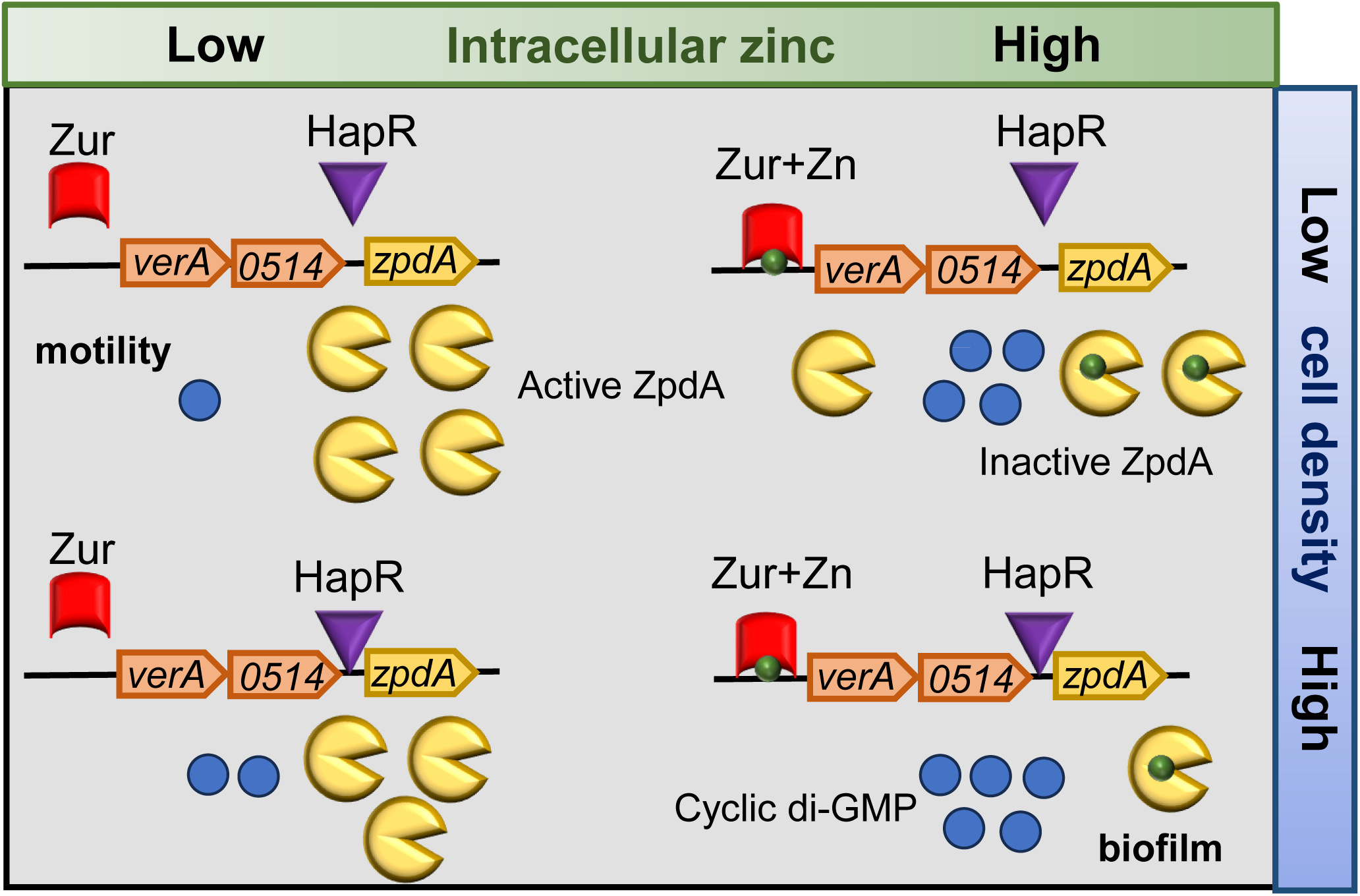
Model describing regulation of ZpdA by Zn^2+^ and quorum sensing. Under very low concentrations of Zn^2+^ and at low cell density, repression by Zur and HapR are relieved resulting in high ZpdA active protein favoring a motile phenotype. At higher Zn^2+^ concentrations, Zur represses the upstream promoter; however, at low cell density *zpdA* is expressed and the protein could be active or inactive depending on the intracellular Zn^2+^ levels. At high Zn^2+^ concentrations and at high cell density, Zur and HapR represses the expression of *zpdA* gene and ZpdA protein will be inactive due to the presence of high Zn^2+^, potentially favoring biofilm formation.

Environmental signals that regulate cdG levels are poorly understood. Bile acids, temperature, spermidine, oxygen and cell density has been show to regulate cdG levels ^14^. Many of the HD-GYP and EAL domains containing PDEs were shown to be bound by metals to regulate their activity *in vitro*^19^. In *V. cholerae* VC0681 binds to Fe^2+^ and Ca^2+^, VieA binds to Mn^2+^, and VC0395 binds to Ca^2+^ for their activity^20,21,38^. Heme and Zn^2+^ was shown to inhibit the activity of the CdpA and VieA PDE respectively in *V. cholerae*^20,39^. However, regulation of PDEs by metals has not been studied in intact bacteria and thus it is not clear whether metals function as signals.

In this study, we demonstrate that extracellular Zn^2+^ serves as a signal modulating the activity of ZpdA at both transcriptional and post-translational levels (Fig. 6). We show ZpdA is active only in the presence of Mn^2+^ and to a lesser extent Mg^2+^ *in vitro* and that Zn^2+^ addition inhibits this activity (Fig. 5C and S3C). Recently it was shown that Mn^2+^ increases the PDE activity of *Pseudomonas aeruginosa* by increasing the activity of RbdA PDE^39^; however, possible inhibition by Zn^2+^ was not addressed. Mn^2+^ or Mg^2+^ binding to EAL domains were proposed to bridge water molecule forming hydroxide ion causing nucleophilic attack on the phosphorus atom of cdG enabling the formation of pGpG^40,41^. The presence of Zn^2+^ could inhibit this nucleophilic attack and hence the PDE activity. The conserved metal binding residues that binds to Mn^2+^ or Mg^2+^ present in other EAL PDEs^40,41^ is also present in the AlphaFold model of ZpdA, hence it is possible that ZpdA could modulate its activity via binding Mn^2+^ or Mg^2+^. Further studies are required to understand how Zn^2+^ might bind and or inhibit it (Fig. S5). In *V. cholerae*, two other PDEs, CdgJ (VC0137) and VC1851, and two PDEs from *Geobacter* and *Bacillus,* share similar domain structures to ZpdA, with N-terminal EAL domains and C-terminal HDOD domains (Fig. S6). Like ZpdA, these EAL enzymes may also modulate their PDE activity through Mn^2+^ and Zn^2+^, in response to environmental cues: we are currently exploring these possibilities (Fig. 6).

*Vibrio cholerae* likely encounters a variety of environments that have variable Zn^2+^ availability. For example, our results show chitin contains significantly higher concentration of zinc of when compared to ocean salts and *V. cholerae* is highly proficient in forming biofilms in the presence of chitin (Fig. S2)^26,42^. The human host induces nutritional immunity by sequestering essential metals such as Zn^2+^ and Fe^3+16–18,43,44^. Possibly, in environments with high Zn^2+^ concentrations, such as chitin, *V. cholerae* forms biofilms through the repression of ZpdA and other PDEs. In contrast, during human infections or in estuarine environments, *V. cholerae* may encounter low Zn^2+^, which triggers switching to an enhanced motility state. The gene upstream of *zpdA, vc0514,* might also enable motility as it was predicted to contain HAMP domain along with a methyl-accepting chemotaxis domain. Previous findings have shown *verA* and *aerB* to be involved in aerotaxis^28^. Hence, the presence of this operon in the VSP-2 island might be important for switching between motile and sessile phenotype by expression of *verA, vc0514* and *zpdA* in low and high Zn^2+^ environments (Fig. 6). The Zn^2+^ responsive transcriptional control of this operon, modulated by QS, coupled with a Zn^2+^-responsive post-translational control of cdG levels, could be critical for environmental persistence. This behavior is reminiscent of uropathogenic *Escherichia coli*, where changes in Zn^2+^ availability provide cues that allow the organisms to persist and adapt to growth in diverse ecological niches^45^. Our finding that *V. cholerae* imports up to 3.5-fold more Mn when switched from zinc-replete to zinc limiting growth conditions, taken together with the finding that addition of Mn^2+^ stimulates PDE activity of purified ZpdA, suggest both Zn^2+^ and Mn^2+^ physiology influence the ability of *V. cholerae* to rapidly switch between motile to sessile states in response to a wide range of environmental stresses.

## Materials and Methods

### Media and growth conditions

All bacterial strains and plasmids used in this study are listed in Supplementary Table S1. *V. cholerae* strains used in this study were derived from N16961 unless otherwise indicated as E7946, C6706 Str2. *V. cholerae* strains were grown in LB, Minimal media (1X M9 minimal salts, 2 mM MgS0_4_, 0.1 mM CaCl_2_, 0.4% glucose), or chitin at 35°C with shaking. Chitin slurry is made as described previously with 8 g of crab chitin powder (Sigma-Aldrich) mixed in 150 ml of 0.5x Instant Ocean water (7 g of instant ocean sea salts (Aquarium Systems, Inc.)/liter of water)^46^. Minimal media was prepared with ultrapure Milli-Q water to minimize metal contamination. ZnSO_4_ was used at the concentration of 10 µM. Cultures were supplemented with appropriate antibiotics when needed. Antibiotics were used at the following concentrations: ampicillin (100 μg/mL), kanamycin (100 μg/mL), polymyxin B (10U/mL), streptomycin (2500 μg/mL). P_tac_ inducible plasmids were induced with 1 mM IPTG.

### Plasmid and strain construction

Plasmids and oligonucleotides used in this study are listed in Supplementary Table S2 and S3. Deletion strains were constructed using pKAS32 suicide vector^47^. For deletions, 700 bp upstream and downstream homologous region were cloned using the KpnI and SacI restriction sites in pKAS32 using Gibson assembly. The plasmids were electroporated into *E. coli* S-17 (λ Pir) strain and conjugated into *V. cholerae* strains. P_tac_ inducible overexpression plasmids were constructed by Gibson assembly of inserts amplified by PCR along with linearized pEVS143 plasmid using EcoRI and BamHI. Plasmids and oligonucleotides used in this study are listed in Supplementary Table S2 and S3. Deletion strains were constructed using pKAS32 suicide vector^47^. For deletions, 700 bp upstream and downstream homologous region were cloned using the KpnI and SacI restriction sites in pKAS32 using Gibson assembly. The plasmids were electroporated into *E. coli* S-17 (λ Pir) strain and conjugated into *V. cholerae* strains. P_tac_ inducible overexpression plasmids were constructed by Gibson assembly of inserts amplified by PCR along with linearized pEVS143 plasmid using EcoRI and BamHI. For protein expression, pET28MBP vector^48^ was amplified using Phusion DNA polymerase (NEB) in GC buffer with primers oMJF001 and oMJF002. PCR product was treated with DpnI and used as the template for a secondary amplification using primers oMJF001 and oMJF172 to add a 3’ StrepII tag to the cloning site. *zpdA* was amplified from gDNA using Q5 polymerase (NEB) with primers oMJF173 and oMJF174. Vector and insert PCR products were mixed 1:1 and transformed into chemically competent DH5α E. coli following the FastCloning procedure^49^.

### Motility assay

For motility assay, the strains were grown in solid media containing 0.375% agar with antibiotics and IPTG as needed. Pipette tips were used to stab the plates from an overnight culture. The plates were inverted and incubated overnight at 35°C in a humidity chamber and imaged using gel imager. The area of motility was analyzed using Hough Circle transform package from the UCB Vision Sciences plugin in Fiji ^50^.

### Biofilm assay

Biofilm formation was determined using MBEC assay as described previously^51^. Briefly, 1:1000 dilution of overnight culture was inoculated in 150 ml of media and incubated overnight at 35° in a humidity chamber. The MBEC lid was washed with PBS for 5 min to remove non-adherent cells. Following washing, the MBEC lid was transferred to 96-well black plate (Perkin Elmer) containing 150 ml of 40% (v/v) Back-titer Glo (Promega) diluted in PBS and incubated for 10 min at room temperature. Following incubation, luminescence was measured using EnVision Multilabel Plate Reader (PerkenElmer).

### Promoter-lux analysis

Overnight cultures were diluted 1:1000 in LB containing pAAR6 encoding the promoter region of *zpdA* in PBBR-lux. The cells were grown with shaking at 35°C in tubes. 150 µl cultures were pipetted into 96 well plates at 2 hours (for low cell density) and 22 hours (for high cell density). The luminescence and OD_600_ was measured using EnVision Multilabel Plate Reader (PerkenElmer).

### Intracellular cdG measurement

The intracellular cdG was quantified as described^23^. Briefly, 1 ml of cultures were spun down and the supernatant was removed. The cell pellets were resuspended in 100 µl cold nucleotide extraction solution (40% acetonitrile, 40% Methanol, 0.1% Formic acid, and 19.9% Water) and incubated at -20°C for 20 minutes to release the nucleotides. The samples were centrifuged, and the supernatant was transferred to a fresh tube and dried under vacuum. The dried samples were resuspended in 100 µl HPLC grade water and analyzed by LC-MS/MS. The concentration of cdG was determined by interpolating with known cdG standards. cdG levels were normalized to total intracellular volume to determine the molarity.

### Protein purification

For MBP-ZpdA and EDTA MBP-ZpdA protein purification, expression plasmid was transformed into BL21(DE3) Star *E. coli* (Invitrogen). Overnight culture was used to inoculate 1 L of complete autoinduction media^52^ with 100 µg/mL kanamycin^52^. Cultures were shaken at 120 RPM for 6 hours at 37°C followed by 16 hours at 18°C. Cells pellets were harvested, washed with PBS, and were frozen at -80°C. To purify the proteins, cell pellets were thawed in an ice-water bath and resuspended in the following lysis buffer: 25 mM HEPES KOH at pH 7.5, 5% glycerol, 300 mM NaCl, 2.5 mM 2-mercaptoethanol, 5 mM EDTA and 0.5% CHAPS along with protease inhibitors (Compete protease inhibitor tablets, Sigma). Resuspended cell slurry was homogenized through a pre-chilled microfluidizer (Microfluidics, Newton, MA). Lysate was clarified by centrifugation at 10,000 RPM in an RC5B centrifuge (Sorvall) an SLA-1500 rotor at 4°C for 30 minutes. Clarified lysate was filtered through a 0.45 µm syringe filter (Millipore, St. Louis, MO) prior to purification. Filtered lysate was loaded onto a 5 mL MBPTrap column (Cytiva, Marlborough, MA) affixed to an AKTA Purifier FPLC (Cytiva) at 2 mL/min using a sample pump in a 4°C refrigerated case. Column was washed with 20 CV of buffer consisting of 25 mM HEPES KOH at pH 7.5, 5% glycerol, 300 mM NaCl, 2.5 mM 2-mercaptoethanol, 5 mM EDTA, followed by 10 washing in 10 CV of EDTA-free buffer. Protein was eluted in column buffer supplemented with 10 mM maltose. Fractions were pooled and dialyzed overnight at 4°C against a storage buffer of 25 mM HEPES KOH at pH 7.5, 10% glycerol, 300 mM NaCl, 5 mM 2-mercaptoethanol. Purified protein was quantified by A_280_ nm using a Nanodrop and the calculated extinction coefficient of 118160 M-1 cm-1. Purified protein was analyzed on SDS-PAGE gel followed by staining with Coomassie. Batches of the protein were also purified without addition of EDTA to the lysis and column buffers. Thus, EDTA was not present in any purified protein used in subsequent enzyme activity assays.

### Protein activity assay

Phosphodiesterase activity of wild-type and EDTA treated protein was determined as described previously with modifications^53^. Protein activity was carried out in 100 µl volume in buffer containing 50 mM Tris HCl at pH 9.5, 50 mM NaCl along with MnCl_2_, ZnCl_2_, MgCl_2,_ CaCl_2,_ FeSO_4_ at the concentration of 1 mM. To this, 0.2 µM enzyme and 2.5 µM cdG were added and incubated at room temperature for 10 minutes. The enzyme was denatured at 90°C for 5 minutes, then cooled and mixed 1:1 with DMHA buffer. The pGpG product was analyzed using LC-MS/MS as described along with the standards of known concentration of pGpG^54^.

### qRT-PCR assays

Overnight cultures were sub-cultured 1:1000 in respective media. For low cell density, OD_600_ of 0.1-0.2 and for HCD, OD_600_ of 2.0-2.5 were used. 1 mL of each replicate was pelleted, and RNA was isolated using TRIzol reagent (Thermo Fisher) according to manufacturer’s instructions. RNA quantity was determined by NanoDrop. 5 µg of RNA was treated with Turbo DNase (Thermo Fisher) and cDNA synthesis was performed using Superscript III reverse transcriptase (Thermo Fisher).

0.78 ng of cDNA was added to 0.625 µM of each primer with 2x SYBR Green Master Mix (Applied Biosystem) in a 25 µL reaction. The reactions were performed in technical triplicates with three biological replicates for each sample, including no reverse transcriptase (no RT) control to monitor for contaminating genomic DNA. qRT–PCR reaction conditions are 95 °C for 20 s, then 40 cycles of 95 °C for 2 s and 60 °C for 30 s in the QuantStudio 3 Real-Time PCR system (Applied Biosystems). *gyrA* was used as an endogenous reference gene for relative quantification (Δ*C*_t_). relative fold expression was calculated using 2^ΔΔCt^method ^55^.

### Inductively coupled plasma mass spectrometry (ICP-MS) analysis

All sample preparation steps were performed using ultrapure Milli-Q water (18.2 MΩ·cm, 25°C) and metal-free polypropylene tubes (15 mL and 50 mL, Labcon). For pellet samples, overnight cultures of *WT* and *ΔznuABC* at 1:1000 were inoculated into 10 mL of minimal media with or without 10 µM ZnSO_4_. The strains were grown for 5 hours after which the cells were washed thrice with 7 mL of 240 mM sucrose solution and centrifuged at 4000 g for 10 min. 100 µL were plated to determine colony forming units at the last wash. After centrifugation, the supernatant was removed, and the pellets were completely dried at 70°C overnight with the conical tube caps loosened. Dried bacterial samples were digested in 150 μL of trace metal-grade nitric acid (Fisher Chemical, Cat. No. A509P212) at 70°C for 2-3 hours or until completely digested. After cooling for 1-2 hours at room temperature, samples were diluted with 4.85 mL of ultrapure water to achieve a final nitric acid concentration of 3%. For powder and culture media samples, the amounts were precisely weighed and subjected to the same digestion protocol described above. Throughout all ICP-MS sample processing steps, empty tubes and samples were weighed to ensure accurate sample volume determination for subsequent concentration analysis.

Elemental quantification was performed on an Agilent 8900 Triple Quadrupole ICP-MS system equipped with an SPS 4 Autosampler, integrated sample introduction system (ISIS), x-lens configuration, and micromist nebulizer. The instrument was tuned using a tuning solution (Agilent, 5185-5959). Gas mode-specific tuning was performed for both helium kinetic energy discrimination (KED) and oxygen reaction modes. An internal standardization approach was implemented using the ISIS valve system with a 200 ppb multi-element solution, containing Bi, In, ^6^Li, Sc, Tb, and Y (IV-ICPMS-71D, Inorganic Ventures). The following isotopes were monitored for comprehensive elemental analysis: ^23^Na, ^24^Mg, ^39^K, ^44^Ca, ^31^P^16^O, ^32^S^16^O, ^52^Cr, ^55^Mn, ^56^Fe, ^57^Fe, ^58^Ni, ^59^Co, ^60^Ni, ^63^Cu, ^64^Zn, ^65^Cu, ^66^Zn. Internal standardization was achieved using ^45^Sc and ^89^Y. Calibration standards were prepared by serial dilution of a 10 ppm multi-element stock solution (IV-65024, Inorganic Ventures). A comprehensive 15-point calibration curve was established with standards from 0.02 to 800 ppb in 3% nitric acid matrix. Elemental concentrations in experimental samples were calculated by interpolation from these calibration curves using Mass Hunter software (Agilent).

## Acknowledgements

We thank Tobias Dörr, Cornell University, for providing N16961, N16961 *Δzur*, N1696*1 Δvc0515*, N16961 *ΔznuABC* strains used in this study. We thank Geoffrey Severin for the construction of the pLJW01 plasmid. Elemental analysis was performed at the Michigan State University Quantitative Bio Element Analysis and Mapping (QBEAM) Center and the National Research Resource for Quantitative Mapping in the life Sciences (QE-Map), generously supported by the Office of the Director, National Institutes of Health and the National Institute for General Medical Sciences (NIGMS) under grant number P41 GM135018. We thank Lijun Chen, Anthony Schilmiller, and Cassandra Johnny of MSU Mass Spectrometry Core for their suggestions and technical support. This work is supported by NIH grants R35 GM139537 and R01 AI158433 to C.M.W., as well as grants R01GM038784 and R01GM115848 to T.V.O.

---

**Fig. S1:** (A) Intracellular cdG levels in wild-type, *Δzur, ΔzpdA* and *Δzur ΔzpdA* for cells grown in minimal media. For comparison, wild-type and *ΔzpdA* quantifications from Figure 1F are repeated. (B) Quantification of motility by measuring the colony area on LB plates for wild-type, *Δzur, ΔzpdA* and *Δzur ΔzpdA* cells. For comparison wild-type and *ΔzpdA* quantifications from Figure 1G are repeated. (C) Quantification of biofilm expressed as relative luminescence units in wild-type, *Δzur, ΔzpdA* and *Δzur ΔzpdA.* Mean and standard deviation of three biological replicates is indicated. Statistical significance was determined by one-way ANOVA and post-hoc Tukey test (* = p < 0.05, *** = p < 0.005, **** = p < 0.0005).

**Fig. S2:** Inductively Coupled Plasma Mass Spectrometry (ICP-MS) quantification of Zn content of growth media: M9-minimal salt, LB powder, ocean salt, shrimp and crab chitin. The average of three independent replicates is indicated.

**Fig. S3:** (A) Relative expression of *zpdA* in E7646 strain at low cell density (LCD) (OD_600_ – 0.1) and high cell density (HCD) (OD_600_ 1.5-2) in LB and chitin media. Mean and standard deviation of three biological replicates is indicated. Statistical significance was determined by t-test (* = p < 0.05, *** = p < 0.005, **** = p < 0.0005). (B) cdG binding RFP biosensor readout normalized to OD_600_ from overexpression of *zpdA* and empty vector in minimal media and LB. For comparison, *zpdA* overexpression and empty vector in minimal media values from Figure 1B. Mean and standard deviation of three biological replicates is indicated. Statistical significance was determined by t-test (* = p < 0.05, *** = p < 0.005, **** = p < 0.0005). (C) Concentration of pGpG measured by LC-MS/MS in wild-type and EDTA-treated protein with cdG substrate in the presence of 1 mM MgCl_2_, 1 mM MnCl_2_, 1mM ZnCl_2_, 1mM FeSO_4_, 1mM CaCl_2_ along with no metal and no substrate controls. Mean and standard deviation of three replicates is indicated. Statistical significance was determined by one-way ANOVA and post-hoc Tukey test (* = p < 0.05, *** = p < 0.005, **** = p < 0.0005).

**Fig. S4:** Comparison of VSP-2 island genes of different El Tor strains are shown. *vc0513 (verA), vc0514, vc0515 (zpdA)* genes are represented in red, other genes are represented in green. Filled and unfilled box represents the presence of gene and absence of gene respectively.

**Fig. S5:** AlphaFold-3 predicted structure of ZpdA protein (AF-Q9KUK3-F1-v4) showing the metal-binding site and domain organization. The structure is displayed highlighting the N-terminal EAL domain and C-terminal HDOD domain arrangement, with two M²⁺ ion (indicates any divalent cation which is predicted to bind to the same site) binding simulated through the AlphaFold Server. Red boxes indicate magnified views of the proposed metal binding site. Conserved residues (E-N-E-E-D-K-E) determined to be involved in metal binding in other EAL proteins are indicated (E90, N68, D118, D119, E166, and K139 shown as green sticks) in proximity to two metal ions (purple spheres), with black dashed lines indicating atoms within 3.0Å distance. The ELL motif residues (E21, L22 and L23, shown in yellow sticks) are positioned near the metal-binding site, with E21 participating in metal coordination.

**Fig. S6:** PANTHER tree viewer (Protein Analysis Through Evolutionary Relationships) of ZpdA protein. Orange circles represent duplications, blue circles represents horizontal gene transfers, green circles represents speciation node, and blue square represents expanded subfamily change node. Blue bars represents EAL domains and pink bars represents HDOD domains of Pfam domain classification.

## Supplementary tables

**Table S1-.**
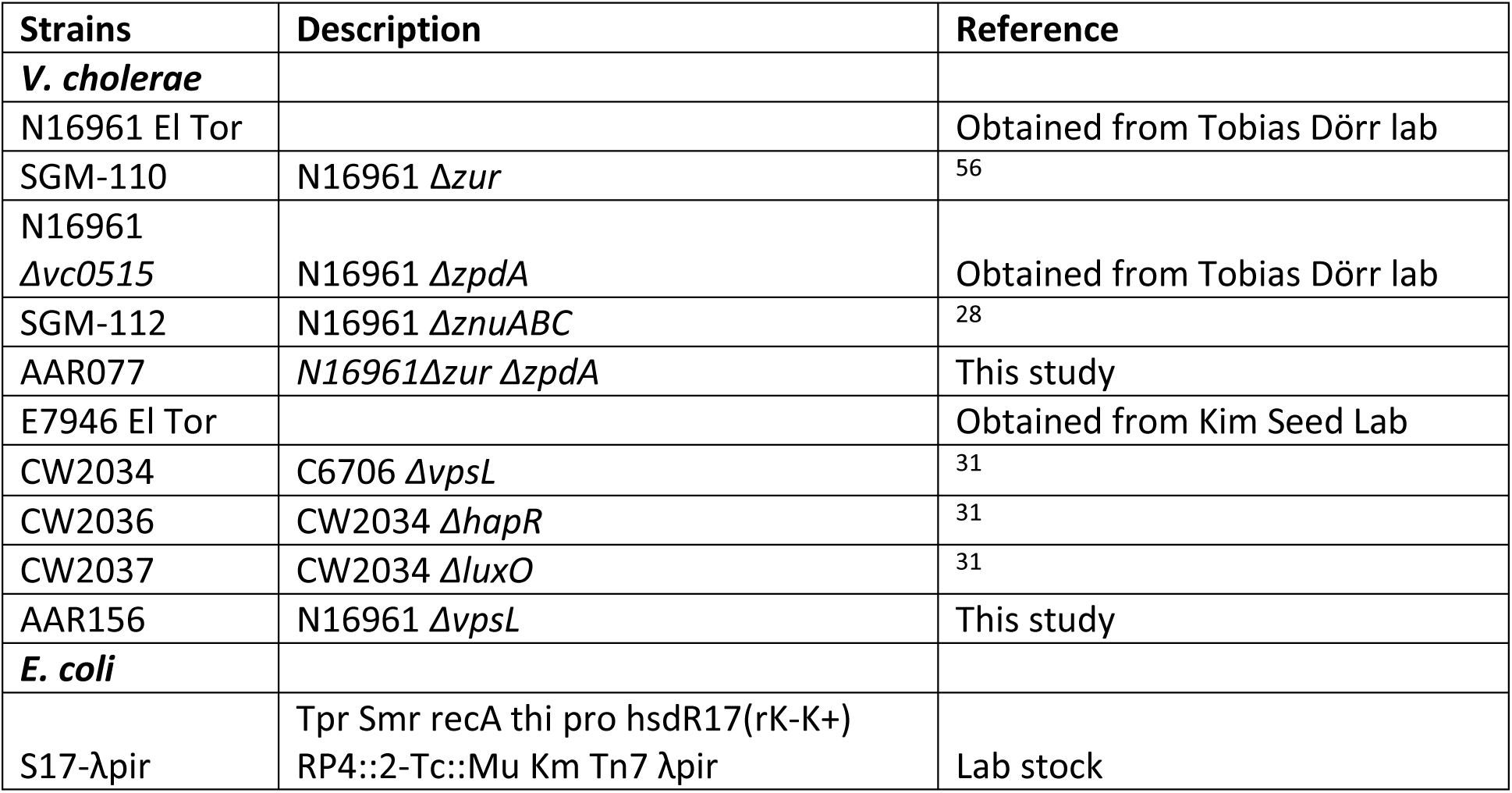
List of strains used in this study.

| Strains | Description | Reference |
| --- | --- | --- |
| *V. cholerae* |  |  |
| N16961 El Tor |  | Obtained from Tobias Dörr lab |
| SGM-110 | N16961 Δzur | 56 |
| N16961 Δvc0515 | N16961 ΔzpdA | Obtained from Tobias Dörr lab |
| SGM-112 | N16961 ΔznuABC | 28 |
| AAR077 | N16961Δzur ΔzpdA | This study |
| E7946 El Tor |  | Obtained from Kim Seed Lab |
| CW2034 | C6706 ΔvpsL | 31 |
| CW2036 | CW2034 ΔhapR | 31 |
| CW2037 | CW2034 ΔluxO | 31 |
| AAR156 | N16961 ΔvpsL | This study |
| *E. coli* |  |  |
| S17-λpir | Tpr Smr recA thi pro hsdR17(rK-K+) RP4::2-Tc::Mu Km Tn7 λpir | Lab stock |

**Table S2-.** List of plasmids used in this study.

| Plasmid | Description | Reference |
| --- | --- | --- |
| pKAS32 | λpir dependent suicide vector, AmpR | 47 |
| pEVS143 | *pTac* overexpression vector, KanR | 57 |
| pBBR-lux | promoterless pBBR1 plasmid with lux genes of *V. harveyi*, CamR | 58 |
| pSLS13 | *hapR* cloned in pEVS143, KanR | 59 |
| pEVS141 | Promoterless pEVS143 vector control, KanR | 60 |
| pVC1353 | *vc1353* in pEVS143, KanR | 61 |
| pCMW121 | *vc1086* in pEVS143, KanR | 31 |
| pRP0122_*Pbe_amcyan_Bc3-4_turborfp* | cdG biosensor plasmid, AmpR | 24 |
| pLJW01 | *zpdA* in pEVS143, KanR | This study |
| pET28-MBP-ST | vector for protein expression, KanR | 48 |
| pAAR14 | *zpdA*<sup>AAA</sup> in pEVS143, KanR | This study |
| pAAR6 | *Pvc0515* in pBBR-lux, CamR | This study |
| pAAR19 | *zpdA* in in pET28-MBP-ST, KanR | This study |

**Table S3-.**
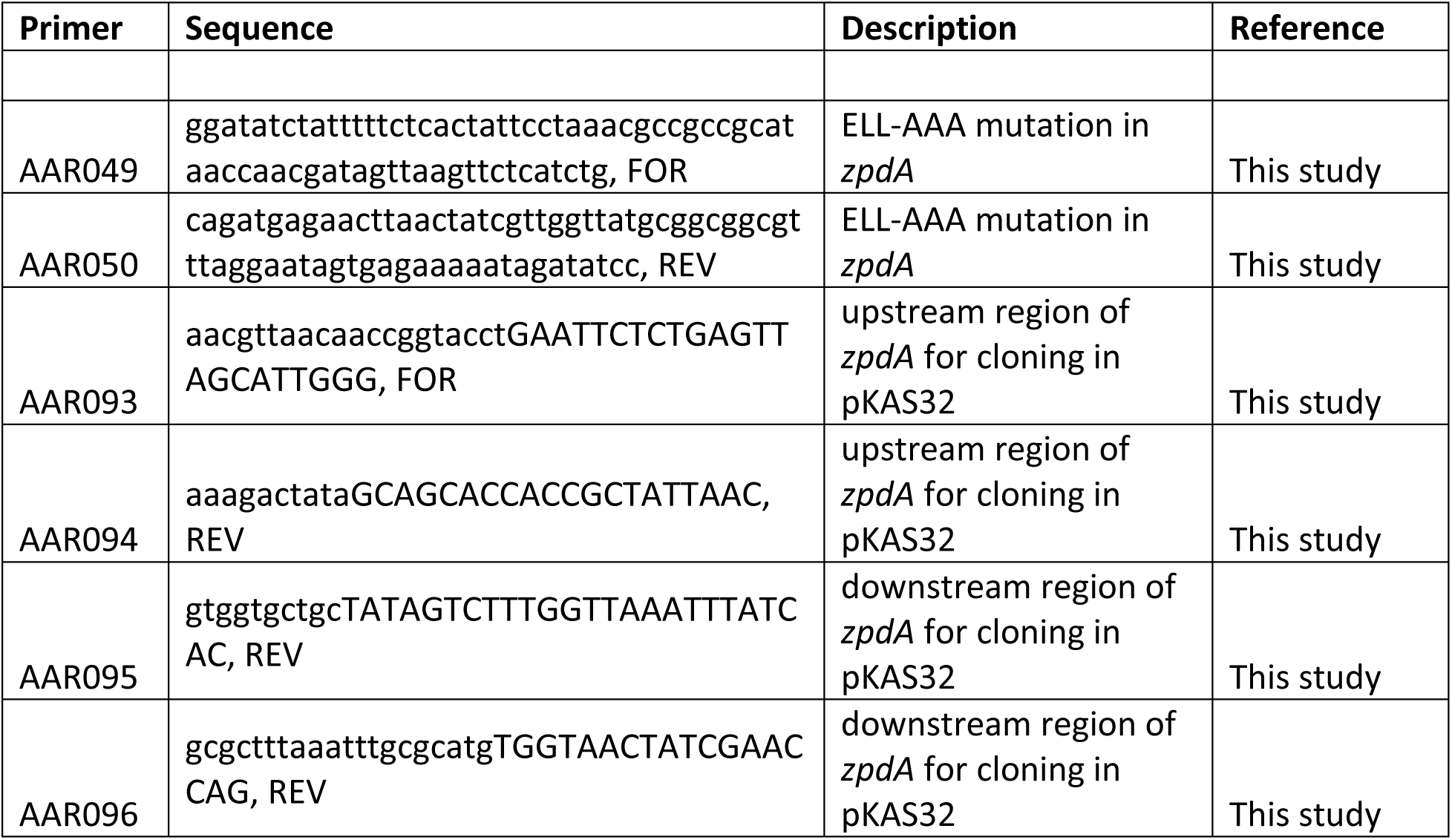
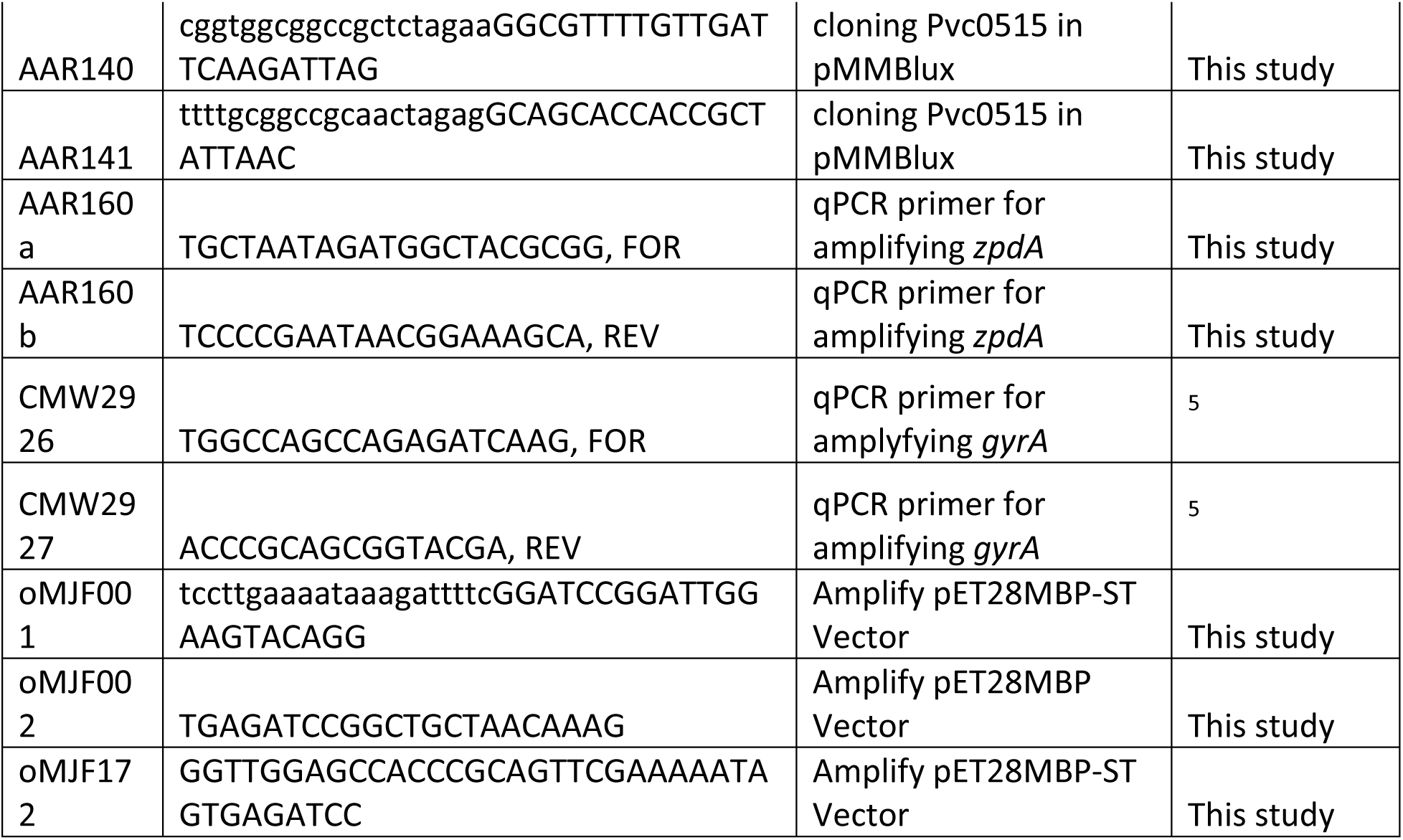
List of oligonucleotides used in this study.

